# A playback paradox? Nathusius’ bats, *Pipistrellus nathusii*, bypass mating and fueling opportunities on migratory transit flights

**DOI:** 10.1101/2022.04.02.486838

**Authors:** Lara C. Marggraf, Oliver Lindecke, Christian C. Voigt, Gunārs Pētersons, Silke L. Voigt-Heucke

## Abstract

In late summer, migratory bats of the temperate zone face the challenge of accomplishing two energy-demanding tasks almost at the same time: migration and mating. Both require information and involve search efforts, such as localizing prey or finding potential mates. In non-migrating bat species, playback studies showed that listening to vocalizations of other bats, both con-and heterospecifics, may help a recipient bat to find foraging patches and mating sites. However, we are still unaware of the degree to which migrating bats depend on con- or heterospecific vocalizations for identifying potential feeding or mating opportunities during nightly transit flights. Here, we investigated the vocal responses of Nathusius’ pipistrelle bats, *Pipistrellus nathusii*, to simulated feeding and courtship aggregations at a coastal migration corridor. We presented migrating bats either feeding buzzes or courtship calls of their own or a heterospecific migratory species, the common noctule, *Nyctalus noctula*. We expected that during migratory transit flights, simulated feeding opportunities would be particularly attractive to bats, as well as simulated mating opportunities which at the same time indicate suitable roosts for a stopover. However, we found that the echolocation call activity of *P. nathusii* decreased during the playback of conspecific feeding buzzes and courtship calls, yet the call activity remained unaffected when heterospecific call types were broadcast. Our results therefore suggest that while on migratory transits, *P. nathusii* circumnavigate conspecific feeding and mating aggregations, possibly to save time or to reduce the risks associated with social interactions. This avoidance behavior could be a result of optimization strategies by *P. nathusii* when performing long-distance migratory flights.

## Introduction

Animals living in temperate zones are exposed to drastic variations in environmental conditions due to a pronounced climatic seasonality. These fluctuations affect prey abundance and habitat suitability, and as a consequence, many species migrate to more favorable areas (Milner-Gulland et al. 2011). Yet, migration is also an energetically challenging task where easy access to relevant information about profitable resources, e.g. foraging and resting opportunities, may be advantageous or even life-saving (Newton and Brocki 2008, Goodale et al. 2010). In some animals of the temperate zone, e.g. in many species of bats, the timing of migration may also overlap with mating activities. These species are confronted with both the challenge of finding sufficient food for fueling the energy-demanding migratory journey with the search for a suitable mating partner at the same time. Information from the environment and from conspecifics, or even heterospecifics may be key for the optimal decision-making in such dual challenge situations (Schoener 1971, Clark and Mangel 1984, Budaev et al. 2019).

In Europe, Nathusius’ pipistrelle bats (*Pipistrellus nathusii*) and common noctules (*Nyctalus noctula*) move within short time from the familiar locations of their summer area to poorly or even unknown areas (e.g. stopover sites) with temporally and spatially unpredictable availabilities of food and roosts (Hedenström 2009). A recent study showed that *P. nathusii* exhibited high metabolic rates during migratory transit flights, even when flying at an energetically optimal speed (Troxell et al. 2019). To cover the elevated energy demands of transit flights, *P. nathusii* use a ‘mixed-fuel strategy’ based on oxidizing ingested insect proteins and fatty acids from their body reserves (Voigt et al. 2012). Although *P. nathusii* depend on insects as an oxidative fuel for migration, they rarely engage in foraging while flying in a migration corridor (Voigt et al. 2018). Instead, they seem to forage first at nightfall and then launch into the sky to proceed their migration route. However, *P. nathusii* is well known to also engage in courtship and mating activities during their stopovers along the migration routes (Schmidt 1994a,b, Furmankiewicz 2003, Jahelková & Horáček 2011). Thus, both of these energy and time demanding life-history stages, mating and migration, are largely overlapping in *P. nathusii*, and also in some other migratory bat species, such as Soprano pipistrelles (*P. pygmaeus*), common noctules (*N. noctula*) and Leisler’s bats (*N. leisleri*) (Dietz & von Helversen 2009).

A solution to the problem of finding profitable foraging sites, suitable mating partners or a roost for resting could either be active communication with other bats via directed social calls, or passive information transfer, i.e., eavesdropping on foraging or courtship behavior of other bats. Indeed, eavesdropping on echolocation calls has been documented for several bat species (e.g. Barclay 1982, Gillam 2007, Dechmann et al. 2013, Übernickel et al. 2012, Cvikel et al. 2015, Roeleke et al. 2020, also reviewed by Gager 2019). At the same time, listening bats which use vocalizations from other bats for additional information acquisition may save energy because echolocation is energetically costly at high intensities (Currie et al. 2020). This is by extending their own range of perception using other bats as an “array of sensors”, i.e. their calls, to detect distant or clumped insect patches, etc. (Fenton 2003, Jones and Siemers 2011, Cvikel et al. 2015, Roeleke et al. 2020). This is facilitated by characteristic, stereotypic repetitions of echolocation calls, so-called feeding buzzes emitted by hunting bats (Schnitzler & Kalko 2001). Eavesdropping on con- and heterospecific echolocation calls, including feeding buzzes, may also help to avoid situations where competition over limited (prey) resources is high (Roeleke et al. 2018). Additionally, flying bats may also locate suitable resting sites by eavesdropping on inadvertent echolocation calls emitted by roosting bats (Ruczyński et al. 2009). Finally, active information transfer with respect to social vocalizations, such as courtship calls or songs, has also been demonstrated for bats. For example, playback experiments showed that bats use social calls to actively coordinate group-foraging (Wilkinson and Boughman 1998). Further, female bats may use male songs to find potential mates (Knörnschild et al. 2017) and possibly suitable roosts. In summary, both passive and active acoustic information transfers represent a prominent behavior in many bat species. Yet, it is unknown whether either active or passive acoustic information is of relevance in the dual challenge situation of bat migration, when bats might trade potential feeding and social activities with the straight continuation of migratory transit flights.

The purpose of our study was to determine whether or not either form of acoustic information transfers, active or passive, play a role for migratory *P. nathusii* during transit flights. We hypothesized that during migration, bats of this species utilizse eavesdropping on feeding buzzes to localize promising foraging patches (passive information transfer by another forager) and, secondly, that *P. nathusii* listen to courtship vocalizations in order to detect suitable mating partners and roosts for stopovers (active information transfer by conspecifics). We therefore predicted *P. nathusii* to be attracted to feeding buzzes during migratory transit flights and thus demonstrate phonotaxis, and to courtship calls, respectively. Further, we assumed that *P. nathusii* would be more attracted to calls of their own species than to calls of heterospecifics, such as *N. noctula*. However, based on similar energetic challenges during migratory transit flights, and the fact that both species are insectivorous, we would expect *P. nathusii* to respond positively, i.e. with increased activity due to positive phonotaxis to *N. noctula* calls. In contrast to this, we predicted that *P. nathusii* would not increase activity at the migration corridor when courtship calls of heterospecific *N. noctula* are played back.

This is the first study to elucidate if and how broadcast acoustic information of bat vocalizations is weighed by actively migrating bats, especially when their need of finding suitable foraging patches and mating partners coincide seasonally and a decision is crucial for both survival (optimal migration) and fitness (optimal mating).

## Material and methods

### Study site

We carried out field work next to ‘Pape Bird Ringing Station’ (56.1667, 21.0059; henceforth abbreviated as PBRS) at the Latvian coastline of the Baltic Sea from 12^th^ of August to 3^rd^ of September 2015. This field site lies within a well-known flight corridor for coastal bat migration used, in particular, by *P. nathusii, P. pygmaeus* and *N. noctula* (Pētersons 2004, Lindecke et al. 2019). To the best of our knowledge, PBRS is solely used as migration corridor as we are not aware of any mating roosts and courting males in this area. We conducted all experiments on a small clearing at a dune, 100 meter inland off the Baltic Sea shore. Migrating bats traverse it, flying along the shore from North to South (Lindecke et al. 2015, 2019). Because of strongly directional flights, we expected to never encounter an animal twice at the experimental site. In support of this notion, we have never encountered any recaptures of the same banded individual within one season.

### Stimulus acquisition

In our playback experiment, we used two functional vocalization types and two stimulus species: foraging call sequences and sequences of courtship calls, both of *P. nathusii* (focal species) and *N. noctula* (control) (Fig. 1).

**Fig. 1.**
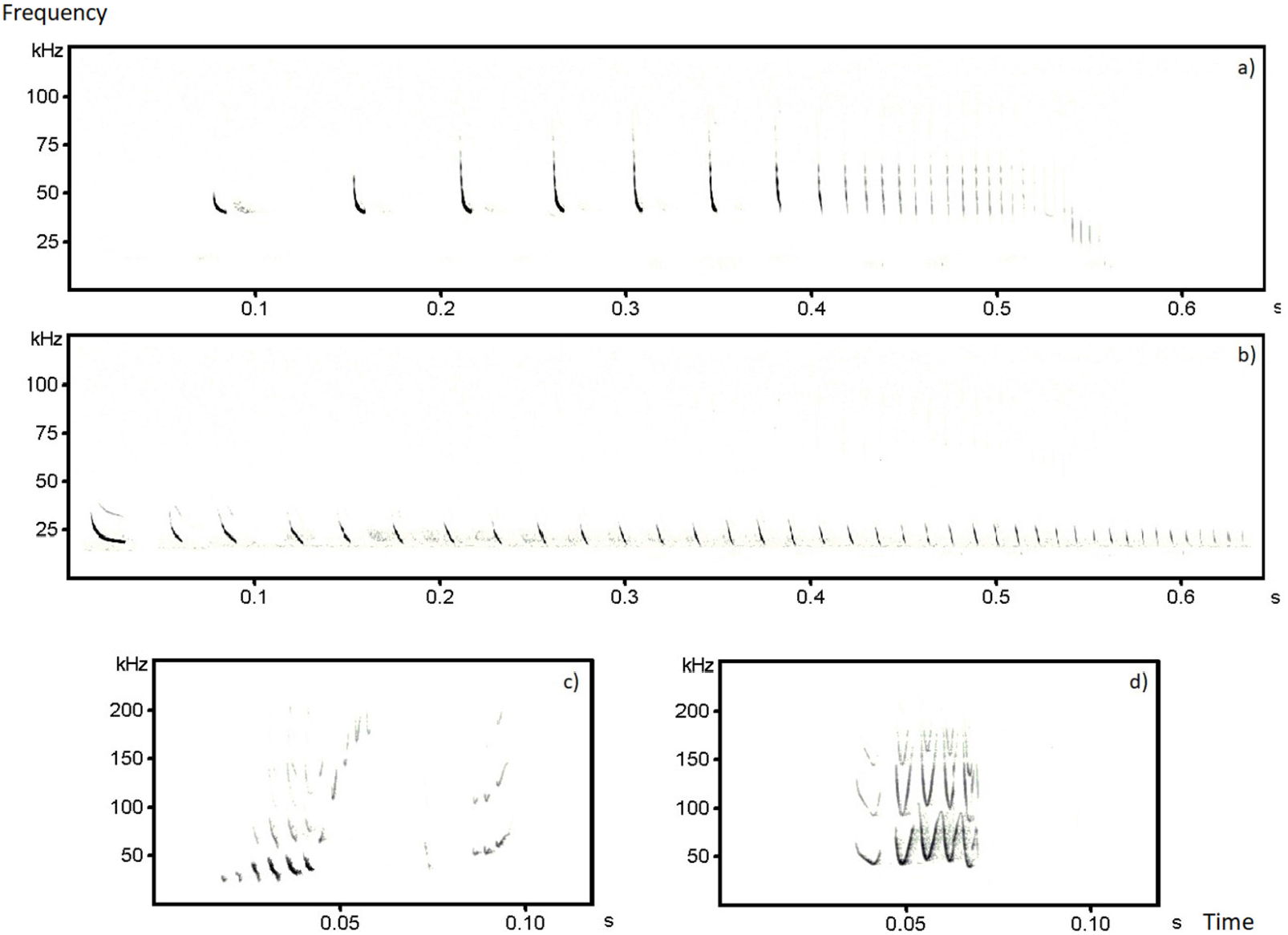
Spectrograms (frequency (kHz) in relation to time (s)) of examples for the stimulus type ‘feeding buzzes’ of *P. nathusii* (a) and *N. noctula* (b). The courtship vocalization of a male *P. nathusii* (c) and the main motif of a male *N. noctula* courtship song (d).

We only chose recordings with a good signal to noise ratio. To create the playbacks of feeding buzzes, we selected sequences from data recorded in the surroundings of Dedelow, Brandenburg, Germany (53.3631, 13.8085) from May to September 2013 and 2014, i.e. from an area in 575 km airline distance southwesterly to our experimental site at PBRS. This sampling region is well within the European mating area of *P. nathusii* (Schmidt 1994a,b) and, in particular, bats passing PBRS may stopover there (ringing data, see e.g. Pētersons 2004). We created files of equal length for every single feeding buzz sequence to about 0.6 s by adding their natural search calls in the beginning. Every single sequence consisted of a search phase, followed by an approach phase and the final buzz phase (Fig. 1a, b). Final playback files with a 1 min duration were created by randomly selecting five 0.6 s sequences which was then replicated in a loop. In total, every 1 min file contained 100 feeding buzzes. This way, we produced 8 individual playback files for each species. For the second vocalization type, the courtship vocalizations, we used files that were also recorded in northeastern Germany during the mating seasons 2010 and 2011 (for detailed information see Voigt-Heucke et al. 2016). For *P. nathusii*, we used vocalizations produced as part of the advertisement song (Fig. 1c) and for *N. noctula* the most common motif of noctule courtship song (Fig. 1d). About 30 individual song motifs per file were randomly pasted together for each species including species-specific characteristics like natural pause lengths between the song motifs. Those sequences were then repeated to obtain a file of 1 min total length. This way, we also obtained 8 individual playback files for each species. We treated final playback files with a high-pass filter at 10 kHz and a low-pass filter at 125 kHz to eliminate background noise. Additionally, peak amplitudes of playback files were separately normalized to 75%. All playback files were created using Avisoft SAS Lab Pro (Avisoft Bioacoustics; Raimund Specht, Berlin, Germany).

### Playback experiment

We placed an ultrasound speaker (ScanSpeak, Avisoft Bioacoustics) and an ultrasound microphone (Avisoft condenser ultrasound microphone CM16, Avisoft Bioacoustics) in 3 m distance to each other to broadcast playback sequences and simultaneously monitor vocal responses of passing bats (Fig. 2). The speaker was placed about 3.5 m above ground; hanging on a dead branch of a pine tree. The microphone was placed in front of the speaker at a height of 1.5 m and orientated upwards. Playbacks were broadcast with an USG Player 116 (Avisoft Bioacoustics, Berlin, Germany), recorded onto RECORDER USGH (Avisoft Bioacoustics, Berlin, Germany) and an ultrasound speaker (TYPE Avisoft Bioacoustics, Berlin, Germany).

**Fig. 2.**
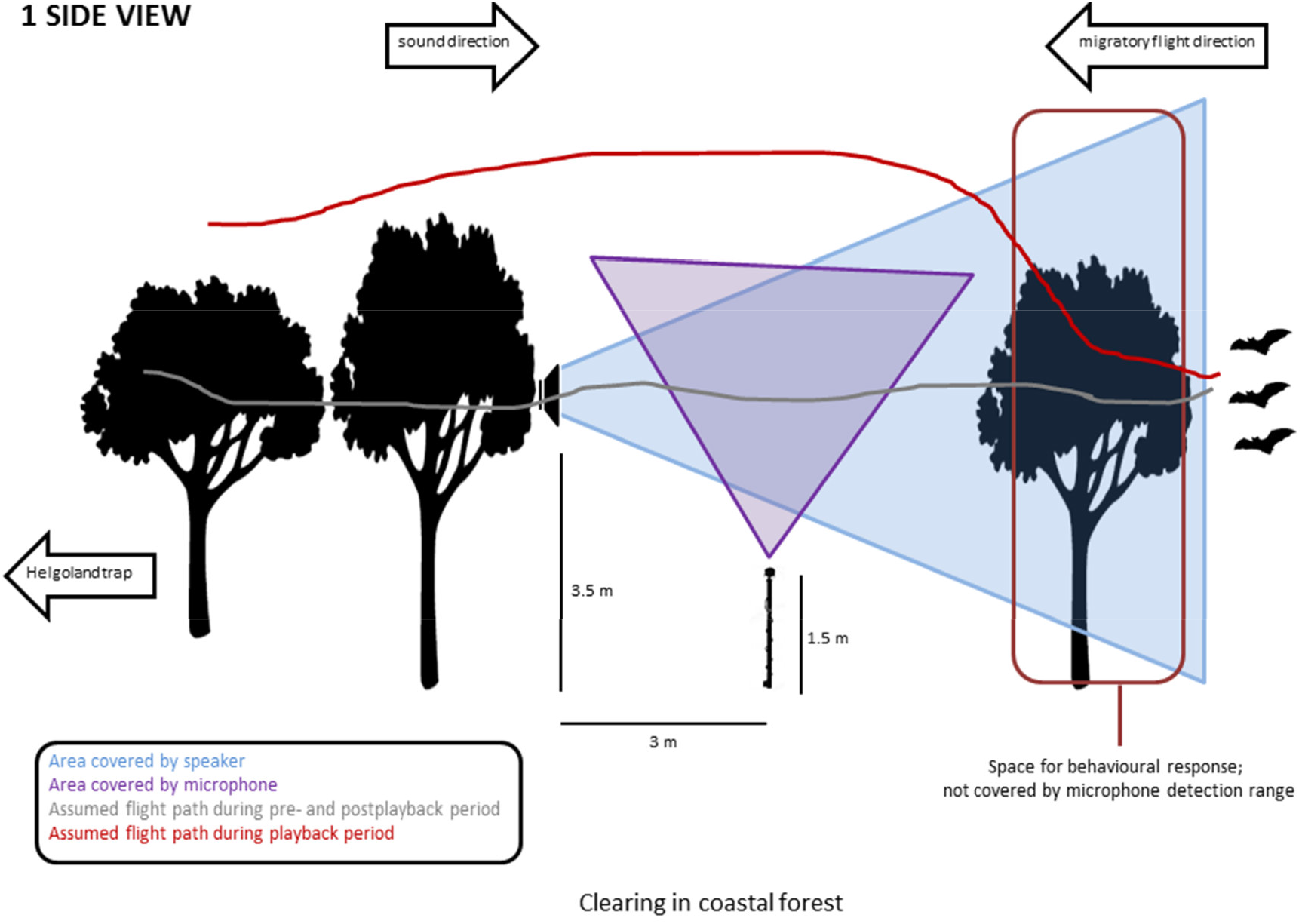
Schematic representation of the experimental set up from side view. The ultrasound speaker was set to the opposite direction of bats on migratory flight. The direction of the microphone was upwards to detect echolocation calls of passing bats, but covered not the whole range of broadcasted signals by the speaker. Before detected by the microphone, bats crossed an area in which they could have decided to change their way before reaching the capture volume of the microphone (leading to the observed reduction in activity).

Filenames of the broadcast playback files were listed simultaneously for differentiation between vocal responses of broadcast playbacks later on. Playback volume was maximized without clipping, resulting in maximum playback amplitudes of 97 ± 2 dB SPL at 0° and 100 cm (mean ± *SD*). Assuming a bat hearing threshold of 10 dB, low playback frequencies (20 kHz) can be audible over 77 m while higher frequencies (50 kHz) that suffer stronger atmospheric attenuation can reach over 25 m at 20°C and 70% relative humidity (L. Jakobsen, pers. comm.; calculations based on Mogensen & Møhl 1979). All recordings were conducted using a 16 bit resolution and a 250 kHz sampling rate. The detection range of the microphone for most common echolocation calls (frequency of maximum energy: 37-42 kHz) of *P. nathusii* was 32 m at 94.2 dB SPL, 0° and 100 cm distance to the speaker. Louder calls of up to 127 dB would be detectable further away, but are still within the range of our playbacks without considering effects of air temperature, relative humidity, position to the bat in relation to the microphone and intensity adjustments by emitting bats according to targets (R. Specht, Avisoft, pers. comm.; see also Barataud 2015 and Adams et al. 2012).

Broadcast playback files consisted of three periods: a 1-min pre-playback period, in which we recorded the baseline for bat activity; a 1-min playback period, in which we presented a playback stimulus and during which we recorded immediate changes to the stimuli; and a 1-min post-playback period, to verify that there was a constant activity of bats passing. We broadcasted two stimulus types: feeding buzzes and courtship vocalisations of two species (resulting in 4 different playback files per playback trial). In each playback phase, one of the four playback files was broadcasted and for each experimental trial, the order of playback stimulus presentation was randomized. In total, one playback trial was 12 min long and started only if we detected calls of *P. nathusii* with our microphone. Each night of the experimental season, we measured wind speed as a proxy for the likelihood of migratory activity approximately 30 minutes after sunset. Playback trials were run subsequently, if wind force was below 8 m/s as migrating bats usually stop flying at high wind speed (Rydell et al. 2010, Cryan et al. 2014, Voigt et al. 2018).

All playback trials were conducted at the same location at PBRS, starting approximately 30 min after sunset. Ideally, playback trials were run throughout the night, except bad weather conditions hindered us from conducting an experiment. The likelihood of presenting a playback to the same individual was negligible as bats continuously migrate towards the South (Lindecke et al. 2015, Voigt et al. 2018) and thus pseudo-replication was avoided. The number of playback trials that we were able to conduct differed between 1 to 15 trials for a given night due to changing weather conditions and general bat activity.

### Analysis of playback recordings

For further analysis, we only included recordings in which vocal activity of *P. nathusii* was present in both the pre-playback and the post-playback period in order to control for a constant activity, i.e. constant bat passings during any experiment. Due to this we post-hoc gathered variable numbers of recordings for each stimulus type. However, this resulted in 99 recordings for playbacks of *P. nathusii* (48 experimental files for feeding buzzes and 51 files for courtship calls, respectively) and 88 recordings for playbacks of *N. noctula* (42 experimental files for feeding buzzes, and 46 files for courtship calls). We counted the number of all echolocation calls (EC) in each of the three periods of a playback experiment to quantify the vocal response of *P. nathusii* to the different stimulus types. For each of these periods, we also counted the number of whole recorded feeding buzzes (FB) and social calls (SC). All acoustical analyses of experimental recordings were performed using Avisoft SAS Lab Pro (Avisoft Bioacoustics) spectrograms (Hamming window, 512 FFT length, 50% overlap).

### Statistical analysis

We tested for normal distribution using Shapiro-Wilk-Tests, which revealed non-normally distributed dataset. To test for differences in vocal responses for each presented stimulus type, we compared the number of EC across time periods (pre-, play-, and post-playback period) using Friedman-Tests. In presence of significant differences, we used Nemenyis Tests as post-hoc tests. All statistical analysis were conducted in R (Version 0.98.1103 – © 2009-2014 RStudio, Inc.). The significance level was set to 5%.

## Results

We conducted playback experiments on 19 nights over the course of three weeks. From 187 experimental playbacks trials, we collected 44,842 calls, consisting of 44,767 EC (99.83%), 63 FB (0.14%), and 12 SC (0.03%). The median number of recorded calls per trial was 11 EC (range: 0-724), 0 FB (range: 0-6), and 0 SC (range: 0-6).

The vocal response of *P. nathusii* to the stimulus types quantified as the number of EC differed among the three playback periods. More precisely, while hearing the playback of conspecific feeding buzzes, we recorded less EC of *P. nathusii* compared to the pre-playback period (Friedman-Test; n = 48, χ2_2_=19.8, p < 0.001; post-hoc Nemenyis test p = 0.0002; Fig. 3) and also compared to the post-playback period (Friedman-Test: n = 48, χ2_2_=19.8, p < 0.001; post-hoc Nemenyis test p = 0.0009; Fig 3). The number of EC decreased by 18.9% between pre-playback and playback period and increased by 17.1% between playback and post-playback period. EC activity of migrating *P. nathusii* did not vary in response to heterospecific feeding buzzes when compared between the periods prior, during and after the presentation of the playback stimulus (Friedman-Test: n = 42, χ2_2_ = 1.29, p = 0.53; Fig. 3).

**Fig. 3.**
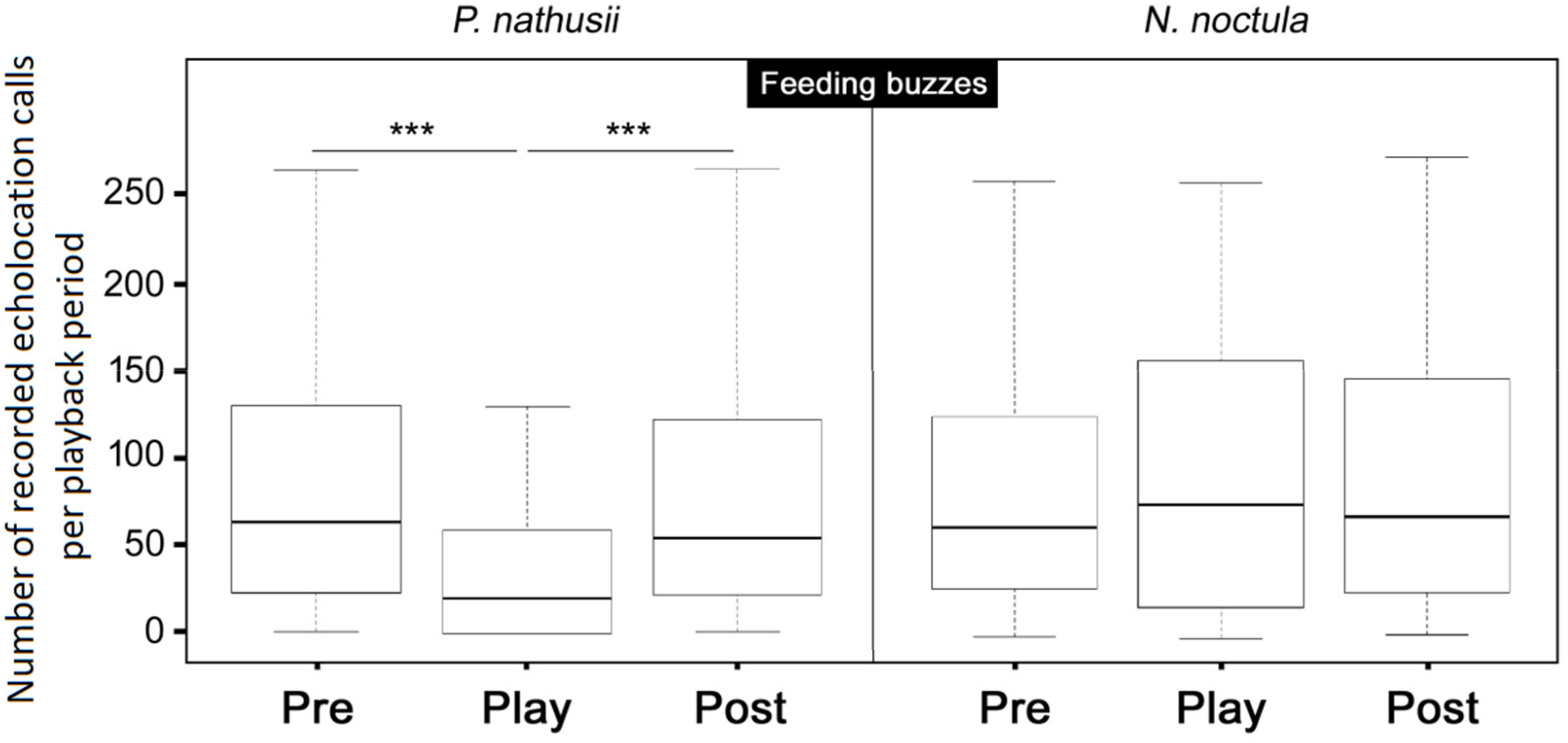
Vocal responses of Nathusius’ bats (*Pipistrellus nathusii*) prior, during and after the playback of feeding buzzes of their own (left graph) and a heterospecific species (*Nyctalus noctula*, right graph). Solid black lines in the center of boxes represent the median, the borders of boxes are 25 and 75 percentiles; whiskers represent the 5 and 95 percentiles. Significant differences between playback periods are indicated by a line associated with an asterisk (*** = p < 0.001).

Similar to the behavioral response in the presentation of feeding buzzes, *P. nathusii* showed less EC activity during the playback of conspecific courtship calls compared to the pre-playback period (Friedman-Test; n = 51, χ2_2_ = 15.96, p < 0.001; post-hoc Nemenyis test p = 0.0007; Fig. 4) and the post-playback period (Friedman-Test: n = 51, χ2_2_ = 15.96, p < 0.001; post-hoc Nemenyis test p = 0.004; Fig. 4). The number of EC decreased by 10.6% between pre-playback and playback period and increased by 11.4% between playback and post-playback period. EC activity of migrating *P. nathusii* did not vary in response to heterospecific courtship calls when compared between the periods prior, during and after the presentation of the playback stimulus (Friedman-Test; n = 46, χ2_2_ = 1.65, p = 0.44; Fig. 4).

**Fig. 4.**
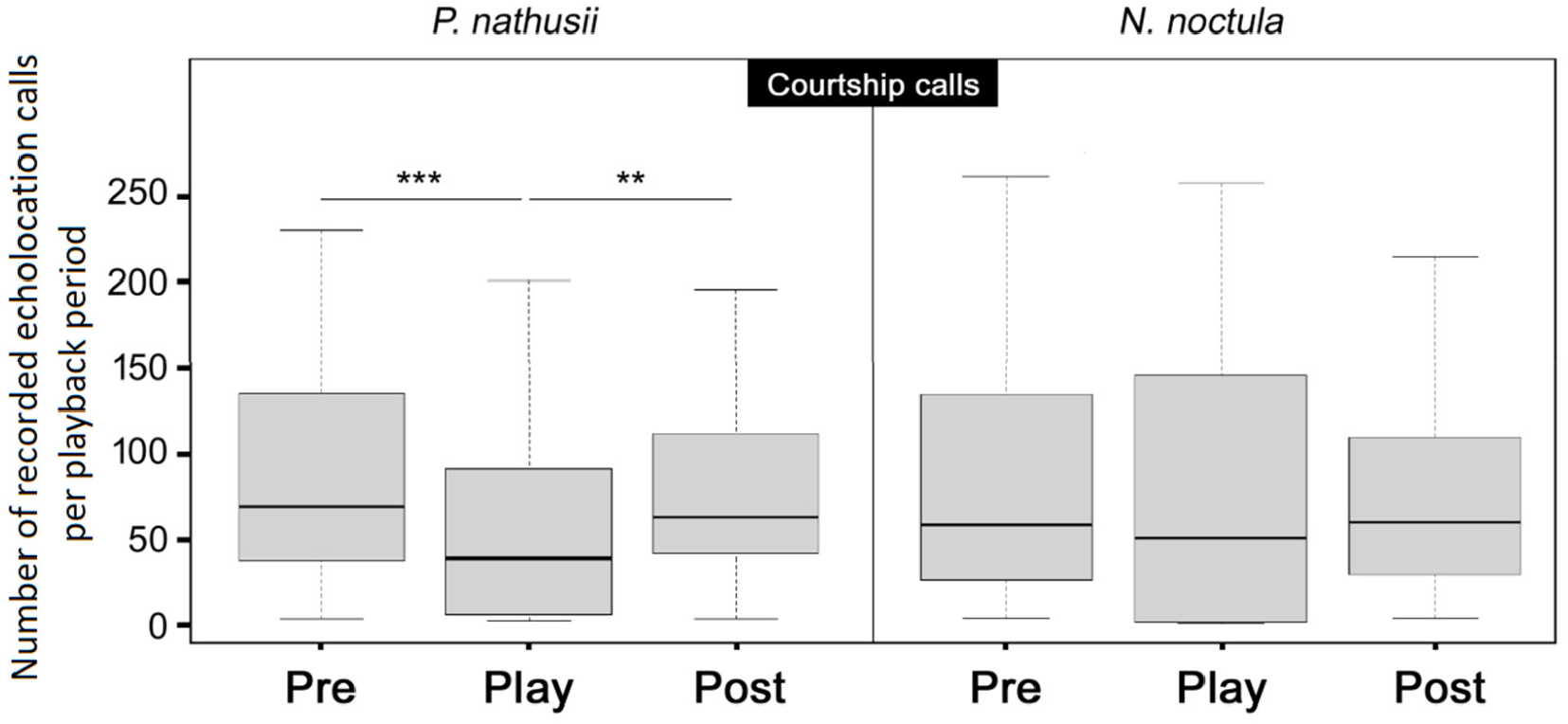
Vocal responses of Nathusius’ bats (*P. nathusii*) prior, during and after playback of courtship calls of their own species (left graph) and a heterospecific species, (*N. noctula*, right graph). Significant differences between playback periods are indicated by a line associated with an asterisk (** = p < 0.01, *** = p < 0.001).

## Discussion

In our study, we investigated the acoustic response of Nathusius’ bats (*P. nathusii*) to simulated feeding and courtship activities of con- and heterospecifics during the annual life-history stage of migration at a major migration corridor for bats in Europe, the coast of the Baltic Sea in Latvia. We expected *P. nathusii* to be attracted to playbacks of FB and to respond to courtship calls of conspecifics during migratory transit flights. We argued that a (simulated) high feeding activity may indicate profitable foraging patches with high insect densities; a valuable resource for migrating bats that encounter high energy demands during migration (Voigt et al. 2016, Costantini et al. 2018, Troxell et al. 2018, Currie et al. 2020). Previous studies demonstrated that some free ranging bat species approach some playbacks of conspecific, and also heterospecific FB. For instance, *P. nathusii* were found to approach loudspeakers that broadcast EC and FB of conspecifics and heterospecifics during late spring and early summer (Dorado-Correa et al. 2013); which is the phase when *P. nathusii* females give birth and wean their young. Approaching behaviour was also found for *P. nathusii* in response to broadcasts of courtship calls in August and September, yet in the non-migratory population of Northern Ireland at the edge of the species distribution range (Russ and Racey 2007). Our study is therefore the first to look at the response behaviour of Nathusius’ bats to conspecific and heterospecific calls during migration.

We found the general EC activity of *P. nathusii* decreased with the playback of conspecific FB and also conspecific courtship calls. Thus, contrary to our predictions, *P. nathusii* appeared to avoid both acoustically simulated feeding and courtship locations. The observed increase in acoustic activity in response to presented stimuli in earlier studies led to the widely accepted conclusion that bats seem to be generally attracted by FB and SC (Russ and Racey 2007, Dechmann et al. 2009; Dorado-Correa et al. 2013; for bat species from other continents and phylogenetic backgrounds see e.g. Gillam 2007, Übernickel et al. 2012). However, none of these studies were conducted in a migratory context and thus, previous studies targeted test animals with different motivations compared to our study. Intriguingly, Roeleke and colleagues (2018) applied our playback files in their study on *N. noctula–P. nathusii* interactions in Germany. They observed that *N. noctula* increased local activity in response to playbacks of *P. nathusii* in early summer when insect densities are high, but reduced their activity in late summer prior to migration onset. Interestingly, in closely related Common pipistrelles, *P. pipistrellus*, Jonker et al. (2010) also found no attraction to the broadcast of conspecific FB and Voigt-Heucke et al. (2016) the same result to the broadcast of SC in studies conducted between August and September. In contrast to *P. nathusii*, however, *P. pipistrellus* seasonally moves over short distances only (∼20 km; Hutterer et al. 2005). Yet, unlike in our experiments, these authors observed no decrease in the acoustic activity of Common pipistrelles (aversion) in response to playbacks. In the situation studied here, we rule out that the aversive behavior of *P. nathusii* in response to conspecific playbacks of FB stimuli might have resulted from an unnatural character of stimuli. Similar stimuli were used in the playback of noctule FB, yet *P. nathusii* did not respond to this heterospecific stimulus, i.e. bats were neither attracted nor repelled by those playbacks. Like in other playback studies with bats, we remain unaware about the exact number of individuals that we tested in our experiment or the sex and age of the recorded bats. Therefore, we cannot make any inferences about the specific behavior of bat individuals, but rather conjecture about the response behavior of *P. nathusii* in general. Keeping these limitations in mind, our study reveals interesting patterns that can be interpreted as aversion to conspecific vocalizations.

Our data revealed a decrease in *P. nathusii* EC activity in response to the broadcast of conspecific FB during the playback period, but no change in EC activity in response to playbacks of heterospecific FB. This finding is contradicting our prediction that during migration eavesdropping on foraging conspecifics might be a strategy to save time and energy. In theory, bats should make use of highly profitable foraging patches that we simulated by the playback of FB. Such acoustic cues should increase the likelihood of finding prey when conspecific bats act as an array of sensors (Gillam 2007, Cvikel et al. 2015, Roeleke et al. 2020). Yet, even though eavesdropping may allow bats to broaden their own range of perception, its use does not necessarily involve advantages only, e.g. bats may need to direct their attention towards conspecifics, and are thus not able to detect prey items at the same time, consequently using more energy for flight maneuvers in order to avoid collision (Amichai et al. 2015). Therefore, we speculate that *P. nathusii* at our study site were not attracted to FB of their own species, because they were anticipating disadvantages from hunting in proximity of unfamiliar conspecifics (Voigt-Heucke et al. 2010). By using a playback of 100 FB/min, we simulated a relatively high feeding activity which might also be interpreted by passing bats as a high density of conspecifics to. Thus, while a high number of FB may indicate a dense cluster of insects on the one side, it could also expose the approaching bat to higher levels of conspecific interference, i.e. aggression on the other side (Racey and Swift 1985). Accordingly, the avoidance of the simulated feeding area may be a strategy to circumnavigate a highly competitive and thus potentially dangerous situation.

We further observed that the EC activity of *P. nathusii* bats decreased in response to the playback of conspecific courtship calls. Most previous studies documented that broadcasting SC will attract target bats or lead to an increase in acoustic activity. For example, in a group cohesion context, Wilkinson and Boughman (1998) demonstrated that social calls of neotropical *Phyllostomus hastatus* attracted conspecifics at roosts and on feeding sites. In a courtship context, some species of bats use calls or even complex songs to attract potential mates. In another neotropical, yet strictly insectivorous bat, *Saccopteryx bilineata*, it was shown that simulating the presence of singing males attracted dispersing females (Knörnschild et al. 2017). But in another case, social vocalizations such as song have also been shown to cause no response in conspecifics of *Tadarida brasiliensis* (Bohn et al. 2013). In our study, we observed a decrease in EC activity in response to the playback of conspecific courtship calls, suggesting that the stimulus elicited an avoidance behavior of the simulated courtship area. In line with our results in *P. nathusii*, Barlow and Jones (1997) found that during the non-mating or migration phase, *P. pipistrellus* reduced their EC activity in response to the broadcasting of conspecific social calls. Barlow and Jones suggested that social calls similar to courtship vocalizations could be used to scare off individuals when used outside of the mating season (Barlow and Jones 1997). Voigt-Heucke et al. (2016) however found that during the late mating season, the playback of con- and heterospecifc social calls did not lead to a change in general EC activity, but a change in the social call rate of wild *P. pipistrellus*. Thus, playback responses to social calls in *Pipistrellus* bats in general seem to depend on the season, and, also on the calling rate with which the playback was constructed. This remains to be tested.

In our study, we were unaware about the sex of the individuals that listened to our playback treatments. In a study on tropical *Saccopteryx bilineata*, Knörnschild and colleagues showed that playback of male song elicited approach flights of mostly subadult females (Knörnschild et al. 2017). Moreover, female *S. bilineata* preferred songs from the local population over songs from foreign locations, demonstrating that song familiarity influences female phonotaxis. Here, we speculate that similar to birds (Kroodsma and Miller 1996), courtship vocalizations could also serve to repel potentially competing males. Accordingly, migrating male bats might have been repelled by conspecific social vocalization because they are motivated to cover distances instead of engaging in territorial encounters that might lead to aggressive encounters. Female *P. nathusii* might as well ignore social vocalizations because social interactions (mating) might prolong their migratory journey. This scenario argues for an avoidance behaviour of migratory bats when conspecific social vocalizations are heard in an otherwise ideal spatio-temporal context (i.e. feeding or mating opportunities in a migration corridor). Interestingly, a playback-study on the function and context of vocalization in a primate species, the mangabey (*Cercocebus atys*) also revealed that test groups moved away from neighboring and unknown calls, but approached those of their own males (Waser 1977). Indeed, Barlow and Jones (1997) found in *P. pipistrellus* that the playback of conspecific SC led to a reduction of acoustic activity when broadcast outside of the mating season. An alternative interpretation to our results comes from another long-distance mammal migrant, Pacific humpback whales (*Megaptera novaeangliae*). Similar to *P. nathusii*, humpback whales also combine mating and migration. In playback experiments, Tyack (1983) observed approaches of male whales to songs and social sounds, but avoidance in females respectively. Female humpback whales may have tried to protect their social group, and in particular the young, by avoiding conflicts, whereas males approached to defend their group. Moreover, playbacks mediated inter-group avoidance in a study on forest monkey, *Cercocebus albiena*, e.g. to circumvent conflicts (Waser 1975). However, it is not known whether *P. nathusii* migrates in large groups and/or with their offspring. Thus, behavior related to group cohesion, protection or an association with offspring may not play a role for migratory *P. nathusii*, and remains speculative.

In conclusion, we found that *P. nathusii* avoided simulated feeding and courtship sites of conspecifics and did not respond to comparatively simulated heterospecific aggregations at a major European bat migration corridor. Our findings argue against a generalised increase of bat activity in response to playbacks of vocalisations of con- or heterospecifics. We therefore conclude advertent or inadvertent information received from calling con- or heterospecifics does not necessarily play a role for *P. nathusii* on migratory transit flights, even though foraging opportunities and mating partners are important in the general context of migration.

## Ethics statement

All procedures performed in this study involving animals were in accordance with the ethical standards of the University of Latvia, Laboratory of Ornithology (license number 10/2015 of the Latvian Nature Conservation Agency) and adhered to the ASAB/ABS Guidelines for the use of animals in research.

## Acknowledgments

We would like to thank Stefanie Zimmer and Olga Heim for allowing us to use their recordings of echolocation and social calls. Furthermore, we are grateful to Oskars Keišs for his support during field work at PBRS, and Kerstin Kraemer and Peter Busse for their support of the work of L.M. We thank Lasse Jakobsen for discussing our data and calculations on the effective range of emitted stimuli.

